# Non-permissive human conventional CD1c^+^ dendritic cells enable *trans*-infection of human primary renal tubular epithelial cells and protect BK polyomavirus from neutralization

**DOI:** 10.1101/2020.09.04.282426

**Authors:** Mathieu Sikorski, Flora Coulon, Cécile Peltier, Cécile Braudeau, Alexandra Garcia, Matthieu Giraud, Karine Renaudin, Christine Kandel-Aznar, Steven Nedellec, Philipe Hulin, Julien Branchereau, Joëlle Véziers, Pauline Gaboriaud, Antoine Touzé, Julien Burlaud-Gaillard, Régis Josien, Dorian McIlroy, Céline Bressollette-Bodin, Franck Halary

**Affiliations:** Nantes Université, Inserm, CHU Nantes, Center for Research in Transplantation and Immunology UMR1064, ITUN, Nantes, France; CHU Nantes, Laboratoire d’Immunologie, CIMNA, Nantes, France; CHU Nantes, Service d’Anatomie et Cytologie Pathologiques, Nantes, France; MicroPicell imaging facility, Structure Fédérative de Recherche Santé François Bonamy - FED 4203/UMS Inserm 016/CNRS 3556, Nantes, France; CHU Nantes, Service d’urologie, France; CHU Nantes, Service de transplantations rénales, France; Plate-forme SC3M, Structure Fédérative de Recherche Santé François Bonamy - FED 4203/UMS Inserm 016/CNRS 3556, Nantes, France; INSERM, UMRS 1229, RMeS, UFR Odontologie, CHU Nantes, Nantes, France; Infectiologie et Santé Publique, UMR INRAE 1282, UFR de Sciences Pharmaceutiques, Université de Tours, France; Département des Microscopies, Plateforme IBiSA de Microscopie Electronique, Université de Tours, France; CHU Nantes, Laboratoire de virologie, Nantes, France

## Abstract

The BK polyomavirus (BKPyV) is a ubiquitous human virus that persists in the renourinary epithelium. Immunosuppression can lead to BKPyV reactivation in the first year post-transplantation in kidney (KTR) and hematopoietic stem cell transplant recipients. In KTR, persistent DNAemia has been correlated to the occurrence of polyomavirus-associated nephropathy (PVAN) that can lead to graft loss if not properly controlled. Based on recent observations that conventional dendritic cells (cDC) specifically infiltrate PVAN lesions, we hypothesized that those cells could play a role in BKPyV infection. We first demonstrated that monocyte-derived DC (MDDC), an *in vitro* model for mDC, captured BKPyV particles through an unconventional GRAF-1 endocytic pathway. Neither BKPyV particles nor BKPyV-infected cells were shown to activate MDDC. Endocytosed virions were efficiently transmitted to permissive cells and shown to be protected from the antibody-mediated neutralization. Finally, we demonstrated that freshly isolated CD1c^+^ mDC from the blood and kidney parenchyma behaved similarly to MDDC thus extending our results to cells of clinical relevance. This study sheds light on a potential unprecedented CD1c^+^ mDC involvement in the BKPyV infection as a promoter of viral spreading.

## Introduction

The BK polyomavirus (BKPyV) is a small non-enveloped DNA virus. Its icosahedral capsid is mainly composed of the major capsid protein VP1(1–3). Its prevalence in the worldwide population ranges from 80 to 90%(4, 5). Asymptomatic primary infection mostly occurs during childhood(6, 7) followed by a persistent infection in the renourinary epithelium(8). Evidence of BKPyV reactivation was mainly reported in kidney and hematopoietic stem cell allografts(9–12) first marked by viral shedding in urine possibly progressing to BKPyV-DNAemia. Persistent BKPyV-DNAemia above 10^4^ DNA copies/ml has been correlated to PVAN (overall 1-5% of KTR)(13–15). To date, BKPyV remains a significant cause of kidney failure(11, 16).

Over the last ten years, anti-BKPyV cellular and humoral immune responses have been investigated demonstrating a prominent role of both specific CD4+ and CD8+ cytotoxic T lymphocytes (CTL), mainly recognizing the large T antigen (LTAg)- and VP1-derived peptides associated with various HLA molecules(17–20). Although anti-BKPyV responses are likely to be protective enough in healthy individuals, only ten percent of those shed virions in urine suggesting a limited impact of escape mechanisms(5). DC are known to orchestrate anti-viral immune responses mainly through their ability to cross-present viral antigens, thus efficiently priming or activating naïve or memory specific T cells respectively(21). To date, anti-polyomavirus (PyV) CTL responses in mice and humans were analyzed on autologous PBMC or DC stimulation using viral peptide pools thus bypassing the requirement for antigen processing, including endocytosis, and presentation by HLA class I molecules(18, 21, 22). Only few studies addressed the ability of PyV to bind to, promote maturation or infect DC. In mice, Drake and colleagues showed that splenic DC are activated following infection by a murine PyV (MuPyV) strain thus allowing them to prime a CTL response(22). Using another experimental setup, Lenz et al demonstrated that although HPV16, a carcinogenic papillomavirus, and bovine PyV virus-like particles (VLP) enabled bone marrow-derived DC maturation, BKPyV or JCPyV VLP did not(23). More recently, hamster PyV (HaPyV)- and Trichodysplasia Spinulosa-associated PyV-derived VLP were shown to moderately activate murine splenocytes(24). Similarly, SV40 was shown to infect and activate MDDC from rhesus macaques(25). Human MDDC were shown to support β-propiolactone-inactivated BKPyV-derived antigen presentation while remaining unresponsive to native BK- and JCPyV particles(26) possibly due to distinct viral antigen processing induced by inactivation(27). Gedvilaite and colleagues also reported that human MDDC were responsive to MuPyV and HaPyV VLP(26). Mostly, DC, although limited to *in vitro* generated cells, seemed to be unresponsive to BK- or JC PyV direct exposure and poorly responsive to BKPyV-derived antigens in KTR and immunocompetent individuals, as recently proposed by Kaur et al(28). The mechanisms behind such DC unresponsiveness remain to be explored regardless of the presence of immunosuppressive drugs.

In the healthy kidney, cDC, including the CD1c^+^ DC subset, are located within the interstitium(29), close to the renal proximal tubular epithelial cells (hRPTEC), a host cell for BKPyV(30). HRPTEC were shown to negatively regulate cDC activation subsequently leading to the retention of cDC in renal tissues as immature cells(31–33) putatively decreasing antigen presentation by DC. Early stage PVAN is marked by a CD1c^+^ cDC infiltrate(34) and mild inflammation(30, 35, 36). Whether cDC play a role in the pathophysiology of the BKPyV infection apart from their ability to trigger and sustain specific immune responses is still unclear.

Here, we demonstrate for the first time that myeloid DC, ie MDDC and freshly isolated CD1c^+^ cDC from the blood and the kidney of healthy donors, but not plasmacytoid DC were capable of capturing BKPyV particles through the CLIC/GEEC endocytic pathway and transmitting them to hRPTEC without getting activated or infected. We also showed that endocytosed BKPyV particles were protected from antibody-mediated neutralization offering to cDC subsets the possibility to participate in BKPyV spreading in the kidney at least in early steps of the reactivation.

## Methods

### Ethic statements

Biopsies from healthy parts of primitive renal carcinoma patients and blood samples from KTR were collected according to institutional guidelines (CPP Ouest authorization, 11/08/2011) and under patients’ informed consent. All samples are conserved in the ITUN bio collection declared at the french Ministère de l’Enseignement Supérieur et de la Recherche under the reference DC-2011-1399 (09/05/2011).

### Cell isolation and culture

Elutriated blood monocytes were obtained from healthy volunteers (DTC cell-sorting facility, CHU Nantes, France) and differentiated into monocyte derived-dendritic cells (MDDC) as described by Sallusto et al(37). Human myeloid CD1c^+^ DC were isolated from blood and kidney by positive immuno-magnetic selection using anti-CD1c/BDCA-1 microbeads according to the manufacturer’s instructions (Miltenyi Biotec, Bergisch Gladbach, Germany) or on a FACS ARIA (BD Biosciences, Franklin Lakes, NJ), respectively. CD1c^+^ DC were recovered from renal cell suspensions of enzymatically digested macroscopically healthy parts of tumor-bearing kidneys (10-15g). Cell purity typically yielded more than 95%. HRPTEC (Sciencell Research Laboratories, Carlsbad, CA) were cultured in complete EpiCM medium (Sciencell Research Laboratories). LNCaP cells (Caliper LifeSciences, Hopkinton, MA) and HEK 293 TT cells (NCI, Frederick, MD) were cultured in RPMI 1640 or DMEM media respectively, both complemented with 2mM L-glutamine and 10% FBS.

### Virus and virus-like particle preparation

The BKPyV Dunlop strain was a kind gift by Dr Christine H Rinaldo (UiT, Norway). The gIa, gIb2 and gIVb1 VP1 expression vectors were kindly provided by Dr Christopher Buck (NCI, USA)(38). Preparation and titration of the Dunlop strain were performed as described elsewhere(39). Virus-like particles (VLP) were purified on an iodixanol gradient(40). VLP physical titers were determined on a qNano device using NP100 nanopores (detection range from 50 to 330 nm) and CPC70 calibration particles (Izon Science Ltd, Oxford, UK). Both viral particles and VLP were labelled with Alexa Fluor®647 protein labelling kit according to manufacturer’s instructions (Molecular Probes, Eugene, OR). Modified-vaccinia Ankara virus (MVA) was kindly provided by Pr Don Diamon (CoH, Los Angeles, CA).

### *Cis* and *Trans*-infection assays

*Cis*- and *trans*-infection experiments were performed as described previously(41, 42). DC and hRPTEC were infected with BKPyV at MOI 0.1 (Dunlop strain). For trans-infection, BKPyV-loaded DC were washed in PBS after two hours at 37°C then put in contact with a subconfluent hRPTEC monolayer. Controls were prepared similarly. After three to seven days post-infection (dpi), LTAg staining was performed to evaluate infection rates as described before (Moriyama and Sorokin, 2009) and imaged on an Axiovert A1 epifluorescence microscope (Carl Zeiss Microscopy GmbH, Germany) or on a Cellomics ArrayScan VTI HCS Reader (Thermo Scientific) for quantification. 25-50 fields, containing 5000-10000 cells were acquired for each well using HCS Studio Cellomics Scan Version 6.5.0 software at various time points pi. VP1 expression was assessed by western blot (ab53977; Abcam) against β-actin (clone C4; Santa Cruz Biotechnology Inc., Dallas, Texas).

### Quantitative RT-PCR analyses

Total RNA was isolated using the TRIzol reagent (Invitrogen) according to the manufacturers’ instructions. Reverse transcription was performed using M-MLV Reverse Transcriptase and random primers following manufacturer's instructions (Invitrogen, USA). Quantitative PCR on reverse transcribed mRNA was performed using Mastermix (Applied Biosystems) or Premix ExTaq 2x (Takara) reagents and the StepOne Plus (Applied Biosystems) or Rotor-Gene (Qiagen) devices. Primers and probe used to detect *LTAg* mRNA were the following: AgT1 5’- ACTCCCACTCTTCTGTTCCATAGG-3’, AgT2 5’-TCATCAGCCTGATTTTGGAACCT-3’ and AGTS 5’-FAM-TTGGCACCTCTGAGCTAC-BHQ1-3’. Expression levels were normalized to GAPDH using the 2-ΔΔ cycle threshold method.

### Gene expression profiling and datasets deposition

BKPyV-mediated cell reprogramming was analysed after 24 hours by 3′digital gene expression (DGE) RNAseq according to Kilens et al.(43). DGE profiles were generated by counting for each sample the number of unique UMIs associated with each RefSeq genes. DESeq 2 was used to normalize expression with the DESeq function. The analysis design used to perform differential expression with DESeq2 between the infected *vs* non-infected conditions took into account the individual DC donors as a confounding variable. Data supporting our results are openly available in the GEO repository under the following ID: GSE154810.

### Flow cytometry analyses

Titrated Alexa Fluor®647 labelled-VLP were used to stain cDC, LNCaP and HEK 293 TT cells at the indicated concentration. VLP attachment was detected by flow cytometry gated on DAPI negative cells. To assess DC activation, cells were incubated for 24 hours with 10^3^ VLP/cell, 10^3^ BKPyV particles/cell or with 100ng/mL LPS and 1µg/mL R848 (Invivogen, San Diego, CA). Antibodies to CD40 (clone 5C3; BD Biosciences), CD80 (clone L307, BD Biosciences), CD83 (clone HB15e, BD Biosciences), CD86 (clone IT2.2, BD Biosciences), CCR6 (clone 11A9, BD Biosciences), CCR7 (clone 3D12, BD Biosciences) and HLA-DR (clone G46-6, BD Biosciences) were used to monitor DC maturation. Whole blood staining was performed on 500µl blood samples from healthy donors with or without Fc fragment receptor blockers (Miltenyi Biotec). Whole blood staining was done with Alexa Fluor®647 labelled-VLP (2.5µg/ml) and cell subsets were discriminated using the following antibody panel: CD45 (Clone J33; Beckman Coulter, Brea, CA), CD11c (Clone BU15; Beckman Coulter), HLA-DR (Clone L243; BD Biosciences), CD123 (Clone 9F5; BD Biosciences) and Lineage (Lin 1; BD Biosciences). FACS analyses were mainly performed on a LSR II flow cytometer (BD Biosciences).

### Fluorescence microscopy

MDDC were distinguished from hRPTEC by DC-SIGN staining (clone DCN46; BD Biosciences) when required. High-resolution confocal microscopy by structured illumination was performed to assess BKPyV entry into MDDC. Cells were incubated with determined VLP concentrations for one hour at 37°C in culture medium, washed and fixed with 3.7% PFA (PFA; Electron Microscopy Sciences, Hatfield, PA). Plasma membrane (PM) was stained with 5μg/mL Alexa Fluor®488-conjugated WGA (Thermo Fisher Scientific). Images were acquired on a Nikon N-SIM microscope with a dedicated oil immersion objective (x100, NA 1.49 Plan Apo). Three dimensional optical sectioning was done respecting Nyquist sampling rate (15 structure illuminations per plane, per channel), and super resolution image reconstruction was performed using Nikon Imaging Software algorithms. BKPyV particle colocalization with CTxB, GRAF-1 and EEA-1 markers was performed as described above with or w/o 2μg/mL Alexa Fluor®555 conjugated CTxB (Thermo Fisher Scientific), and with anti-GRAF1 (4μg/mL; Novus Biological, Littleton, CO) or anti-EEA1 antibody (BD Biosciences) antibodies in 0.1% BSA PBS O/N at 4°C. Nuclei were counterstained with DAPI. Cells were mounted in ProLong™ mounting medium (Thermo Fisher Scientific) and observed on a LSM Nikon A1RSi microscope (Nikon, Tokyo, Japan) at x60 (NA 1.4). 3D reconstruction was done using the Imaris software (Bitplane, Zurich, Switzerland).

### Transmission electron microscopy

MDDC were prepared for transmission electron microscopy as described elsewhere(42). Ultrathin sections were observed on a JEM 1010 microscope (Jeol Europe SAS, Croissy Sur Seine, France). TEM images of BKPyV particle preparations in negative contrast were obtained as described previously(44, 45).

### ELISA

Supernatants from various MDDC cultures were harvested at indicated times and frozen at −80°C until being analyzed. IL-10 and IL-12p70 were quantified in those culture supernatants by ELISA with BD OptEIA™ human IL-10 and IL-12p70 sets following the manufacturer's instructions (BD Bioscience).

### Statistics

Statistical analyses were performed with the PRISM software (GraphPad Software Inc., version 5.04, La Jolla, CA). Almost exclusively one-way ANOVA with multiple comparison tests were performed to assess significance in this study. Exceptionally, correlation and linear regression studies, Mann-Whitney or Friedman tests were also applied to some data sets. *P*-values lower than 0.05 were considered significant.

## Results

### Human monocyte-derived dendritic cells bind BKPyV particles in a dose- and sialic acid-dependent manner

First, we assessed whether MDDC could bind BKPyV particles using fluorescent-labelled genotype Ib2 (gIb2) BKPyV VLP. VLP integrity was checked by negative contrast TEM (Figure 1A). GIb2 VLP binding was then tested with two BKPyV permissive cell types, namely hRPTEC and HEK293TT but also to MDDC and LNCaP, a BKPyV non-permissive prostatic cancer cell line(46) at various VLP/cell ratios. MDDC effectively bound gIb2 VLP in a dose-dependent manner but to a lesser extent compared to hRPTEC or HEK293TT (Figure 1B), and as expected BKPyV particles were unable to attach to LNCaP cells. MDDC were also shown to bind gIa infectious particles (Dunlop strain; (47)) and VLP at comparable levels (Figure 1C). We further demonstrated that genotypes Ia, Ib2 and IVb1 VLP had similar binding properties to MDDC (Figure 1D). Sialic acids decorating *b*-series gangliosides are known as crucial components for BKPyV infection of hRPTEC and HEK293TT(46). Then, we demonstrated that when MDDC are treated with an appropriate neuraminidase, an enzyme known to specifically remove sialic acid moieties from the PM, gIa, gIb2 and gIVb1 VLP binding was strongly impaired (Figure 1E). Altogether, these results clearly established that MDDC could bind BKPyV from the most frequent genotypes in Europe and Asia in a dose and sialic acid-dependent manner.

**Figure 1:**
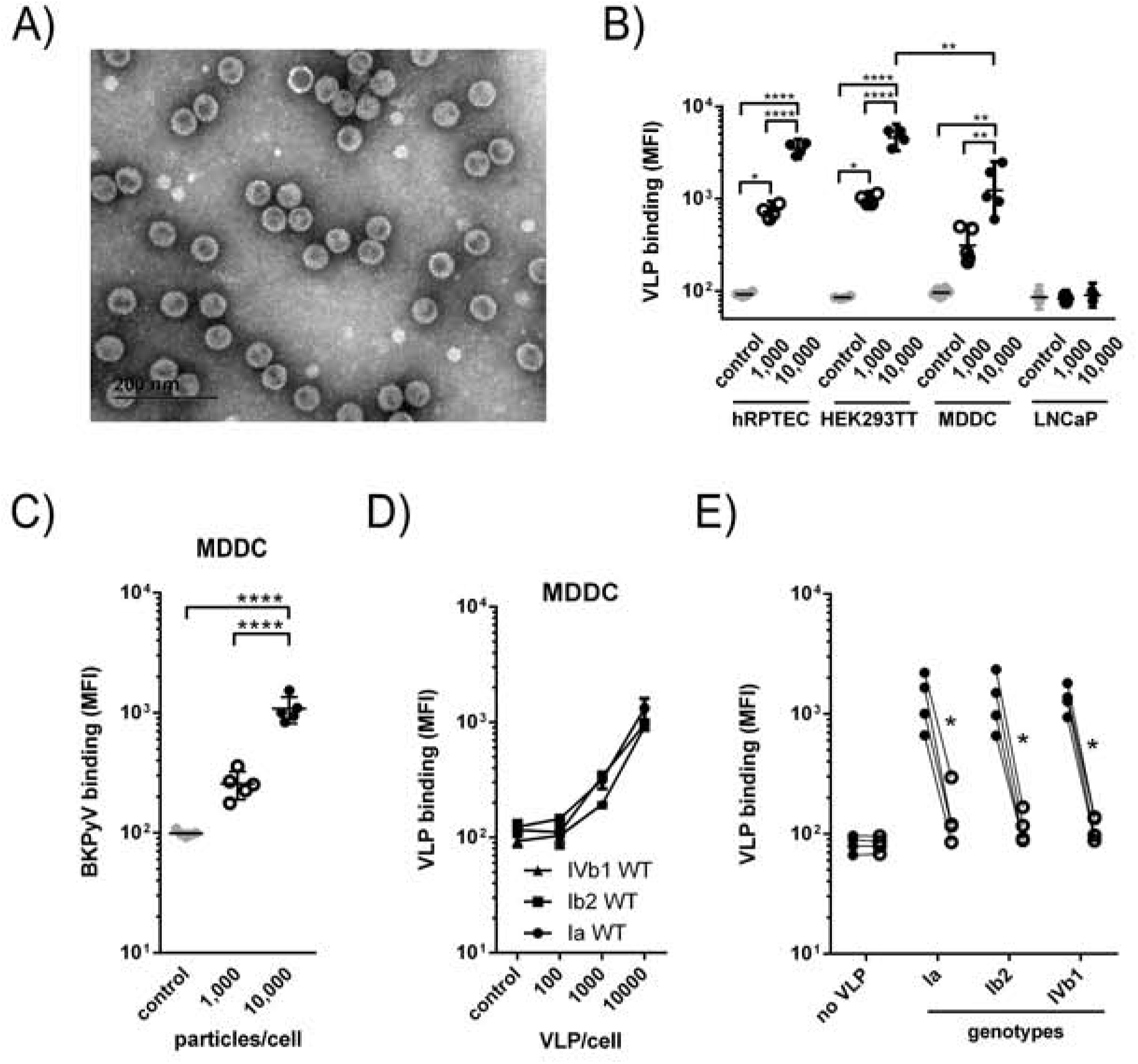
MDDC bind BKPyV particles in a dose- and sialic acid-dependent manner. (A) Negative contrast TEM picture of genotype Ib2 (gIb2) VLP. A 200nm scale bar is represented on the micrograph. (B) Fluorescent-labelled gIb2 VLP binding to hRPTEC, HEK293TT, MDDC and LNCaP (n=3) assessed by flow cytometry. Mean Fluorescence Intensities (MFI) are displayed (n=5; n=3 for LNCaP only). . (C) Alexa Fluor®647-conjugated infectious particles (Dunlop strain) gIb2 VLP binding to MDDC (n=5). (D) Dose dependent binding of genotypes Ia (circle), Ib2 (square) and IVb1 (triangle) VLP to MDDC (n=5). (E) Alexa Fluor®647-conjugated genotypes Ia, Ib2 and IVb1 VLP (10^4^ VLP/cell) binding to MDDC with (empty circles) or w/o (closed circles) treatment with 0.2U/mL neuraminidase from *Clostridium perfringens*, specifically cleaving α(2,3/6/8)-linked sialic acid. Data are represented as MFI ± SEM. Statistically significant results were marked by one or several asterisks according to the level of significance: *=p<0.05, **=p<0.01, ****=p<0.0001; one-way ANOVA with Tukey’s multiple comparison tests.

### BKPyV particles are endocytosed in pleiomorphous tubular and macropinosome-like endosomes in MDDC

Immature MDDC exhibit high endocytic properties for soluble and particulate antigens (37). Therefore, we hypothesized that BKPyV could be endocytosed following attachment to sialic acid residues on PM. High-resolution confocal imaging showed that fluorescent spots representing VLP or virions were found in cytoplasmic structures (Figures 2A and 2B), confirming that MDDC endocytosed BKPyV following surface attachment. VLP were either located in round-shaped or pleiomorphous tubular structures (Figures 2A and 2C). This was confirmed by 3D cell reconstruction (Figure 2D). Then, we performed TEM imaging and confirmed that VLP and virions were mostly internalized after 30 minutes. Indeed VLP were mainly endocytosed into tubular vesicles (40-60nm width) and to a much lower extent in large round-shaped uncoated endosomes (up to approximately 1μm in diameter) by MDDC (Figures 3A, 3B, 3C, 3D3E and 3G). Moreover, these BKPyV-containing tubular vesicles were shown to originate from PM invaginations (Figures 3D and 3E). Some of these vesicles closely resembled sorting endosomes (Figure 3C). Higher magnifications micrographs confirmed that BKPyV virions behaved similarly to VLP (Figure 3F) did not reveal PM curvature upon viral attachment as previously reported for SV40 (Figure 3G; (48)). We concluded that BKPyV was mainly endocytosed into tubular vesicles evoking an uncommon endocytic pathway for viral particles in MDDC.

**Figure 2:**
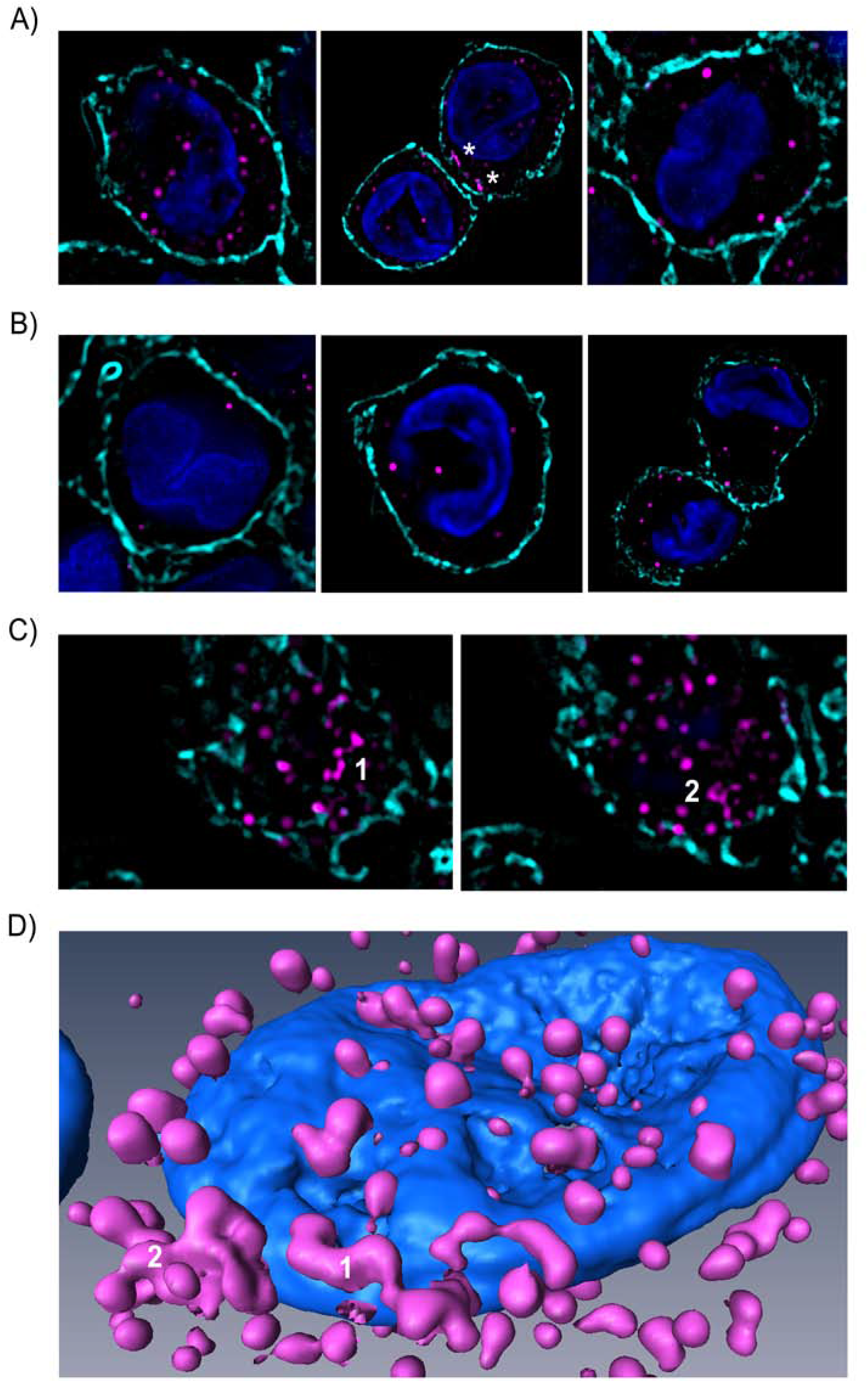
High-resolution confocal images of genotype Ia BKPyV particles endocytosed in MDDC. Three panels showing independent cells that contain intracellular dot-like or amorphous tube-shaped (white asterisk) accumulations of Alexa Fluor®647-conjugated gIa VLP (A) or BKPyV infectious particles (B; Dunlop strain). Images show focal planes extracted from six different cells stacks (104 VLP/cell or 1FFU/cell respectively for VLP and infectious particles; magenta). (C) Two distinct focal planes extracted from the cell stack from which the image in the center of Figure 2A is shown. “1” and “2” indicate the tube-shaped structures marked by asterisks in Figure 2A. (D) Amira 3D reconstruction of the cell represented in Figure 2A (center) showing round-shaped and pleiomorphous tube-shaped intracellular structures containing Alexa Fluor®647-conjugated gIa VLP. Cell membranes were stained with fluorescence-labelled WGA (Alexa Fluor®488 displayed in light blue) and nuclei were counterstained with DAPI. High-resolution confocal images were obtained from the A1 Nikon microscope equipped with a SIM module.

**Figure 3:**
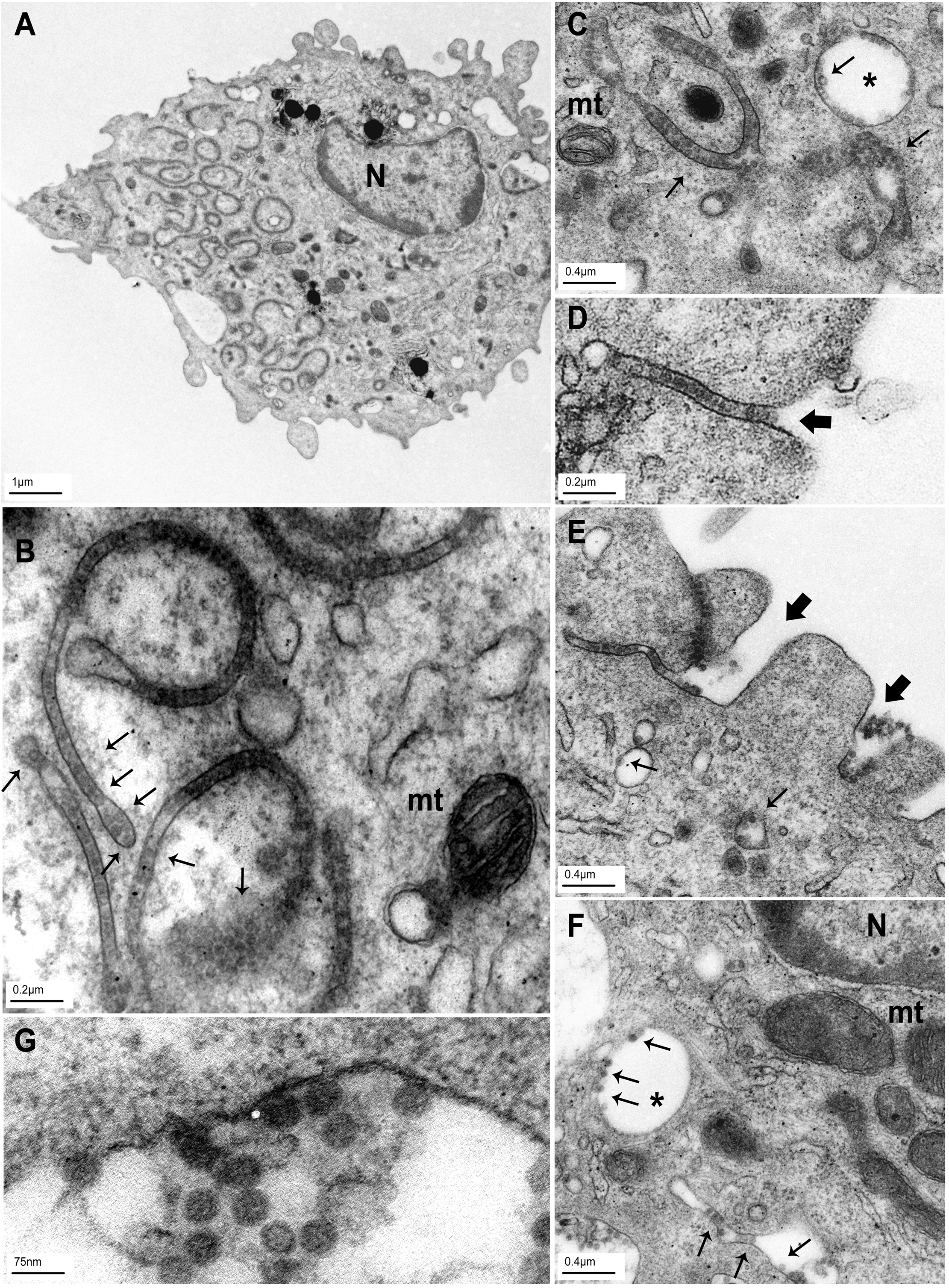
TEM reveals unlabeled BKPyV endocytosis into large round- and tube-shaped vesicles in MDDC. (A) Micrograph showing a general view of a representative MDDC incubated for 30 minutes at 37°C with 10^4^ VLP/cell (gIb2). Noticeably, the cell contains abundant tube-shaped structures. (B, C, D, E and G) Pleiomorphous or large round vesicles containing VLP are shown at a higher magnification (G: x100,000-120,000). (F) This image represents infectious BKPyV particles into macropinosome-like (round-shaped) and tube-shaped vesicles (1FFU/cell). (D) and (E) Micrographs showing VLP internalization from the cell surface into tube-shaped endosomes. Thin and bold arrows indicate particles and tube formation respectively; asterisk indicate large vesicles resembling macropinosomes. N=nucleus; mt=mitochondria. Scale bars are indicated for each micrograph.

### BKPyV colocalizes with GRAF-1+ and cholera toxin B+ compartments in MDDC

To characterize BKPyV containing vesicles in MDDC we used high-resolution confocal microscopy to identify markers co-localizing with BKPyV in MDDC. Early Endosome Antigen-1 (EEA-1), a marker of early endosomes and macropinosomes, was associated with BKPyV in structures with size ranging from 100nm, the detection limit with this technique, to roughly 1μm in diameter (Figures 4A and 4B). The clathrin-independent carriers (CLIC) or GPI-anchored protein-enriched compartments (GEEC) endocytic pathway(49, 50) known to form tubular vesicles has been recently associated with the protein GTPase Regulator Associated with Focal Adhesion Kinase-1 (GRAF1) (51). BKPyV colocalized with GRAF-1 at the PM and in the cytosol (Figure 4C). Cholera toxin B subunit (CTxB) uses GRAF-1 vesicles to enter cells (51) and we observed a partial VLP/CTxB colocalization in MDDC (Figure 4D). Altogether, our results showed for the first time in MDDC a major BKPyV endocytosis into GRAF-1+ and CTxB+ compartments, two hallmarks of the CLIC/GEEC pathway.

**Figure 4:**
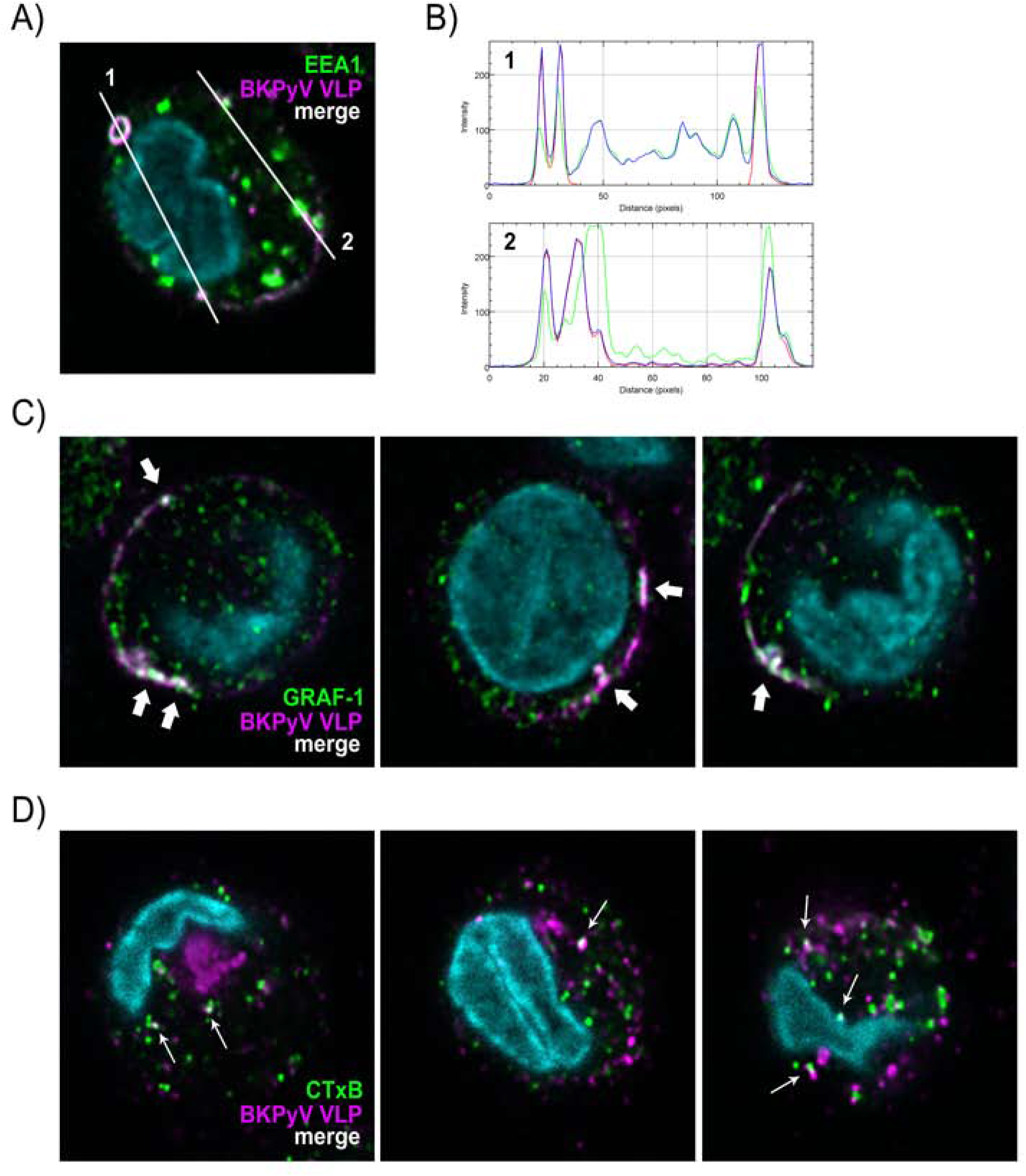
BKPyV particles colocalize with EEA-1, GRAF-1 and CTxB in MDDC revealing an unconventional endocytic pathway. (A) Confocal sections of MDDC incubated for 30 minutes at 37°C with 10^4^ fluorescent VLP/cell (magenta). EEA-1-positive endocytic vesicles were stained after fixation (green). Nuclei were counterstained with DAPI (light blue). The colocalization between VLP and EEA-1 is shown in white. (B) RGB profiles along two measurement lines (1 and 2, showed in Figure 3A) analyzed with ImageJ software. Colocalization is represented by merging blue and red (=magenta) representing VLP and green histograms (1 pixel=88nm). (C) and (D) show respectively colocalization of BKPyV VLP with GRAF-1 (bold white arrows) and Alexa Fluor®555-conjugated cholera toxin subunit B (CTxB; 2μg/mL; thin white arrows). Deconvoluted images are presented. Displayed data are representative of three independent experiments.

### MDDC can transfer virions to renal epithelial cells but are refractory to BKPyV infection

Next, we wondered whether BKPyV-pulsed MDDC, hereafter termed “BKPyV-infected MDDC”, could transfer the virus to a permissive cellular third party in *trans*. Here, we took advantage of a *trans*-infection assay previously set up in our laboratory(41, 52). It assesses the ability of a cell type to capture and transfer virions to permissive cells in its vicinity after removing excess unbound/non-internalized virions. LTAg expression was analyzed in these conditions at defined time points (Figure 5A). Infection of hRPTEC, termed *cis*-infection, was estimated between 10 to 18% in all experiments at seven days pi (Figure 5A). No LTAg was detected in BKPyV-infected MDDC suggesting that BKPyV infection is not initiated in MDDC (Figure 5A). To confirm these results with a more sensitive technique, we analyzed *LTAg* expression by RT-qPCR in a similar experimental design. Quantitative results are shown in Figure 5B. As expected, no LTAg mRNA was detected in BKPyV-infected MDDC whereas the *cis*-infection of hRPTEC or the *trans*-infection conditions displayed high amounts of *LTAg* mRNA. These results were confirmed by assessing the expression of the major capsid protein VP1, a late infection marker (Figure 5C). To confirm the CLIC/GEEC pathway involvement in the BKPyV *trans*-infection process, we finally tested the effect of the ciliobrevin D (CBD), a cytoplasmic dynein inhibitor(53), on MDDC during virus loading. Noticeably, a 50μM dose of CBD significantly decreased *trans*-infection with no measurable MDDC cytotoxicity (Figure 5D). Altogether, these results demonstrated that MDDC, while non-permissive to BKPyV, capture BKPyV virions and can transfer them to permissive cells like hRPTEC in a dynein-dependent manner.

**Figure 5:**
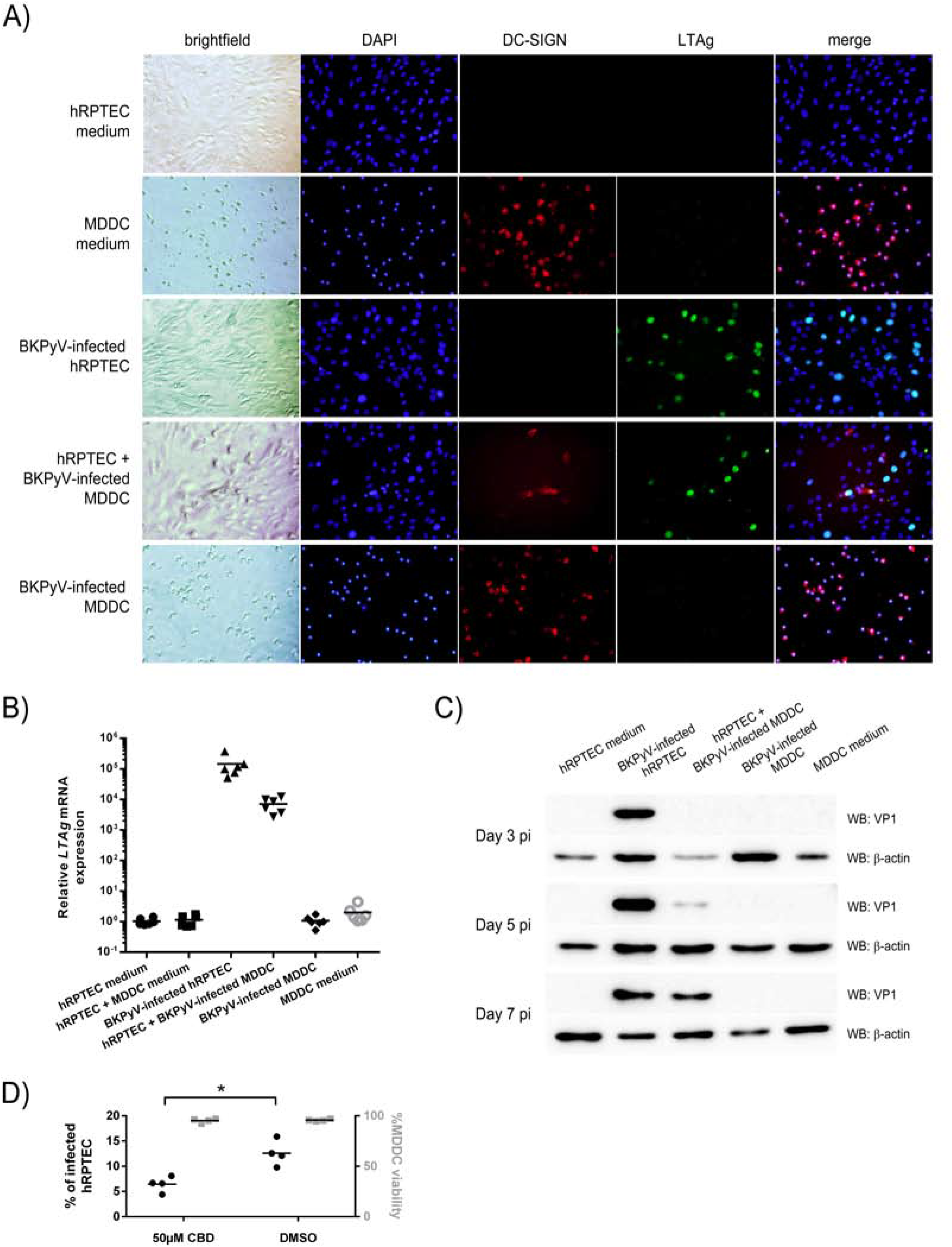
MDDC do not support BKPyV infection but mediate its transmission to primary hRPTEC. (A) Epifluorescence microscope images (x10 magnification) showing large T antigen (LTAg) immunostaining (green) of hRPTEC and/or MDDC in various conditions indicated on top of rows, respectively: hRPTEC alone (medium), MDDC alone (medium), BKPyV-infected hRPTEC (MOI=0.1; approximately 200 particles/cell), non-infected hRPTEC layered with BKPyV-infected MDDC (excess of virus, i.e. unbound virus, was removed by extensive washes after a 2 hour-incubation of MDDC with virus) and BKPyV-infected MDDC (*idem* previous condition). LTAg is revealed at seven days dpi. Brightfield images of the immunostaining are shown in the first column. DC-SIGN (red) is a marker allowing to discriminate MDDC from hRPTEC when necessary. Nuclei were counterstained with DAPI. (B) RT-qPCR data showing the amplification of LTAg mRNA seven dpi in various conditions (similar to those presented in Figure 6A; n=6). Of note, a condition with uninfected MDDC with uninfected hRPTEC has been added here. (C) Western blot analysis of VP1 expression, as a late BKPyV infection event, in cell lysates after three, five, and seven days post-infection. β actin was revealed similarly after membrane stripping as a loading control. Figures 5A and 5C are representative of three independent experiments.

### Human MDDC are neither activated by BKPyV particles nor BKPyV-infected hRPTEC

MDDC can sense danger signals through various pattern-recognition receptors (PRR) including toll-like receptors (TLR) thus leading to MDDC maturation(54, 55). Conflicting results on DC activation by BKPyV in the literature prompted us to ask whether BKPyV attachment would lead to MDDC activation. Twenty-four hour MDDC cultures with VLP or virions were analyzed by flow cytometry to assess the acquisition or up-regulation of known DC maturation markers. A maturation enabling dose of LPS and R848(56), two TLR agonists, and a Modified Vaccinia Ankara (MVA) attenuated poxvirus known to activate MDDC were added as positive controls of maturation when necessary. First, expression of CD86, a sensitive and reliable marker of DC maturation (57), was assessed. Only exposure to TLR agonists or to the Modified Vaccinia Ankara attenuated poxvirus known to activate MDDC induced CD86 upregulation ((58–61); Figure 6A). Accordingly, no IL-12p70, a T helper type 1 cytokine, IL-10 or IL-8 were detected in MDDC culture supernatants cultivated with BKPyV (Figure 6B). Expression of the CD80, CD83, CD40, CCR7 and HLA-DR on MDDC gave consistent results (Figure 6C). Then we hypothesized that MDDC could be activated not by BKPyV particles *per se* but by BKPyV-infected hRPTEC. HRPTEC infection was monitored by RT-qPCR for *LTAg* mRNA expression (data not shown). In that setting, MDDC CD86 expression did not vary upon cultivation with BKPyV-infected cells (Figure 6D). To exclude “under the radar” activation signals, we finally performed a digital RNA sequencing (DGEseq; REF) of BKPyV-infected MDDC compared to non-infected cells. In line with previous experiments, no difference was observed between BKPyV-infected and non-infected MDDC in terms of mRNA profile reprogramming at one dpi (Figure 6E). These results confirmed that MDDC were unresponsive to BKPyV and BKPyV-infected hRPTEC.

**Figure 6:**
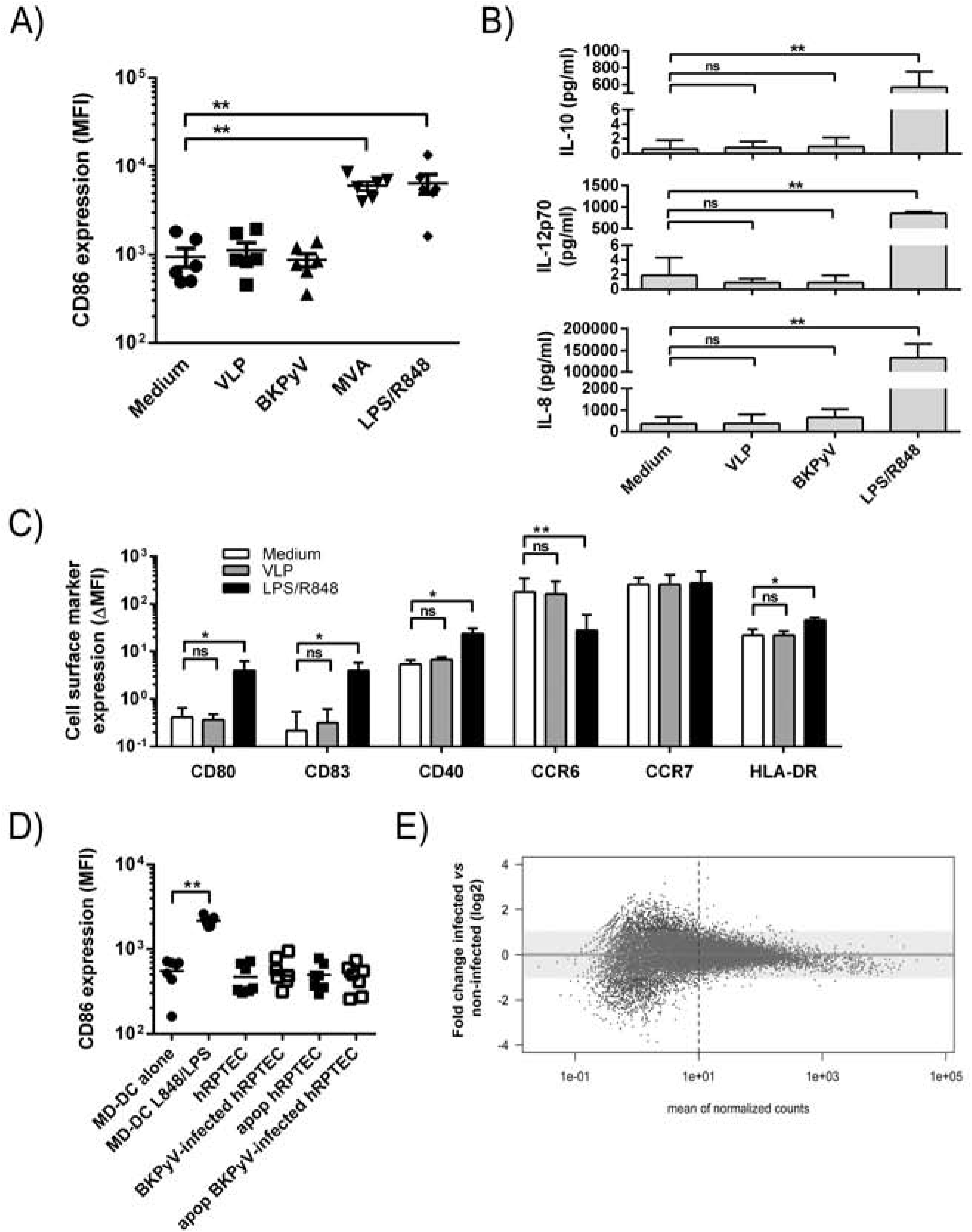
BKPyV virions or BKPyV-infected cells fail to activate MDDC. (A) CD86 cell surface expression was assessed by flow cytometry on immature MDDC alone (circles) or cultured with VLP (squares; 10^3^ particles/cell), BKPyV particles (triangles; 10^3^ particles/cell) MVA (inverted triangles) or a TLR agonist cocktail (diamonds; 100ng/mL LPS and 1μg/mL R848 after 24 hours (n=6). (B) ELISA titration of IL-10, IL-12p70 and IL-8 in the supernatants of untreated or MDDC cultivated with VLP, BKPyV particles or R848/LPS (doses were similar to those employed in Figure 4a). (C) Cell surface expression of CD80, CD83, CD40, CCR6, CCR7 and the HLA-DR on MDDC alone (empty bars) or cultured for 24 hours with VLP (grey bars; 10^3^ particles/cell) or LPS/R848 (black bars). Data are represented as MFI ± SEM. For each MFI, background, i.e. autofluorescence, is subtracted to calculate ΔMFI values displayed in this figure (n=4). (D) Similar to experiments in A. Apop=apoptotic cells. Apoptosis was induced by UVB-irradiation and apoptotic cell fragments were collected by centrifugation and extensive washing in PBS. (E) RNAseq analysis of differentially expressed genes between infected (one dpi) and non-infected MDDC. The dashed line represents a “ten counts per gene” limit above which gene expression is considered as robust. The Y axis represents the Log2 fold change in gene expression. Statistically significant results were marked by one or several asterisks according to the level of significance: ns=non-significant, *=p<0.05, **=p<0.01, ***=p<0.001, ****=p<0.0001; one-way ANOVA with Tukey’s multiple comparison tests.

### Internalized BKPyV is protected from neutralization

Together with cellular immune responses, neutralizing anti-BKPyV antibodies (NAbs) are required to control infection or reactivation in KTR(17, 62–66) and healthy donors(67). Here we wondered whether BKPyV could be protected from neutralization when internalized by MDDC. To address this point, *trans*-infection was performed in the presence of neutralizing and control sera from BKPyV reactivating or non-reactivating KTR respectively. Neutralizing antibodies completely blocked hRPTEC *cis*-infection whereas the control serum had no effect (Figure 7). When virions were pre-incubated with NAbs prior to MDDC loading, a significant loss in *trans*-infection was observed compared to controls. As a conclusion, NAbs were ineffective when used after BKPyV loading of MDDC suggesting virions were protected from neutralization once internalized.

**Figure 7:**
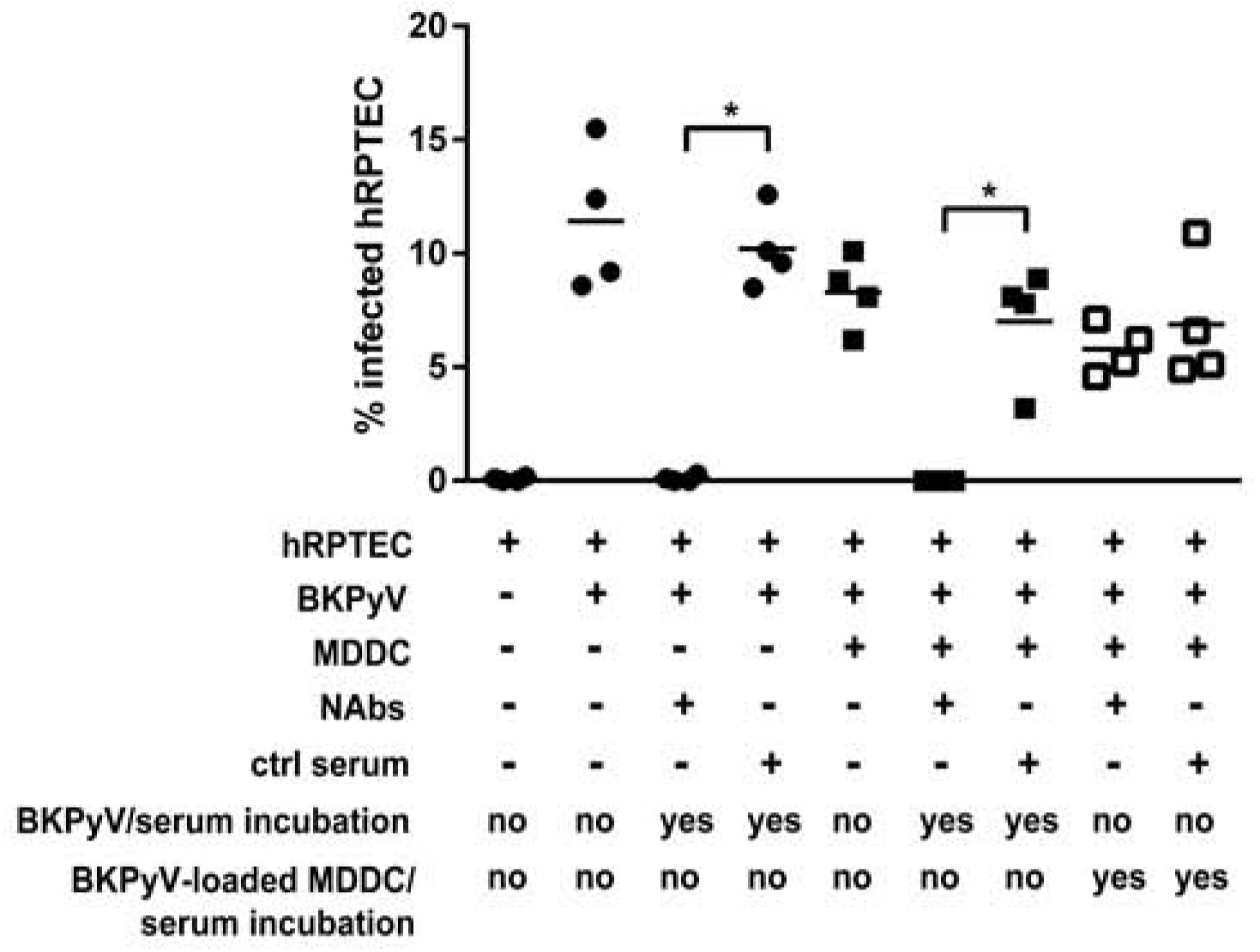
Endocytosed BKPyV particles into MDDC are protected from serum neutralization. HCS automated counting to evaluate percentages of BKPyV-infected hRPTEC (=LTAg+ cells) in various conditions including *cis*- and *trans*-infection experiments but with or w/o sera from a non-controller patient (=control serum) or a controller patient (=neutralizing serum). Neutralizing antibody titers in this serum had been previously determined: between 1/200.000, 1/500.000 and 1/20.000 for genotypes Ia, Ib2 and IVc2 respectively (see the Materials and Methods section). Here, both sera were x1000-diluted. Sera were either added before or after incubation of the BKPyV suspension (Dunlop strain at MOI=0.1) with MDDC. Results represent mean values of the percentage of LTAg+ hRPTEC ± SEM. Statistically significant results are marked by an asterisk; *=p<0.05; one-way ANOVA with Tukey’s multiple comparison tests.

### Blood and kidney CD1c+ cDC display similar BKPyV *trans*-infection abilities and non-permissiveness to MDDC

MDDC were shown to be closely related to inflammatory DC in humans(68, 69) so to ensure our observations were not biased by the DC generation protocol, we first wondered whether cDC, the most abundant tissue and blood DC subset under non-inflammatory conditions, could behave like MDDC. First, VLP were incubated with whole blood of healthy volunteers and VLP staining was further analyzed by flow cytometry on both cDC (CD11c+) and plasmacytoid (pDC; CD123+) DC among HLA-DR+ Lin-cells (Supplemental Figure 1). A significant proportion of cDC bound VLP whereas no binding was detected on pDC (Supplemental Figure 1 and Figure 8A). Importantly, binding to cDC was not affected by Fc receptor blockade suggesting that VLP attachment did not depend on anti-VP1 antibodies in whole blood of healthy donors (Figure 8B). To avoid any interference due to the whole blood environment, CD1c^+^ DC, representing the main myeloid DC subset in blood(70) and kidney(71) were sorted according to the gating strategy displayed in Supplemental Figure 2, incubated with VLP and analyzed by flow cytometry. Blood and kidney CD1c^+^ cDC were clearly capable of binding VLP as well as virions in a dose-dependent manner (Figures 8C and 8D, respectively). Then, we showed that like MDDC, CD1c^+^ cDC were unresponsive to BKPyV particles (Figure 8E). We finally demonstrated that sorted CD1c^+^ cDC enabled BKPyV *trans*-infection to permissive cells while being resistant to infection themselves (Figure 8F). Taken together our results demonstrate that biologically relevant blood and renal CD1c^+^ cDC behave similarly to MDDC with respect to BKPyV infection.

**Figure 8:**
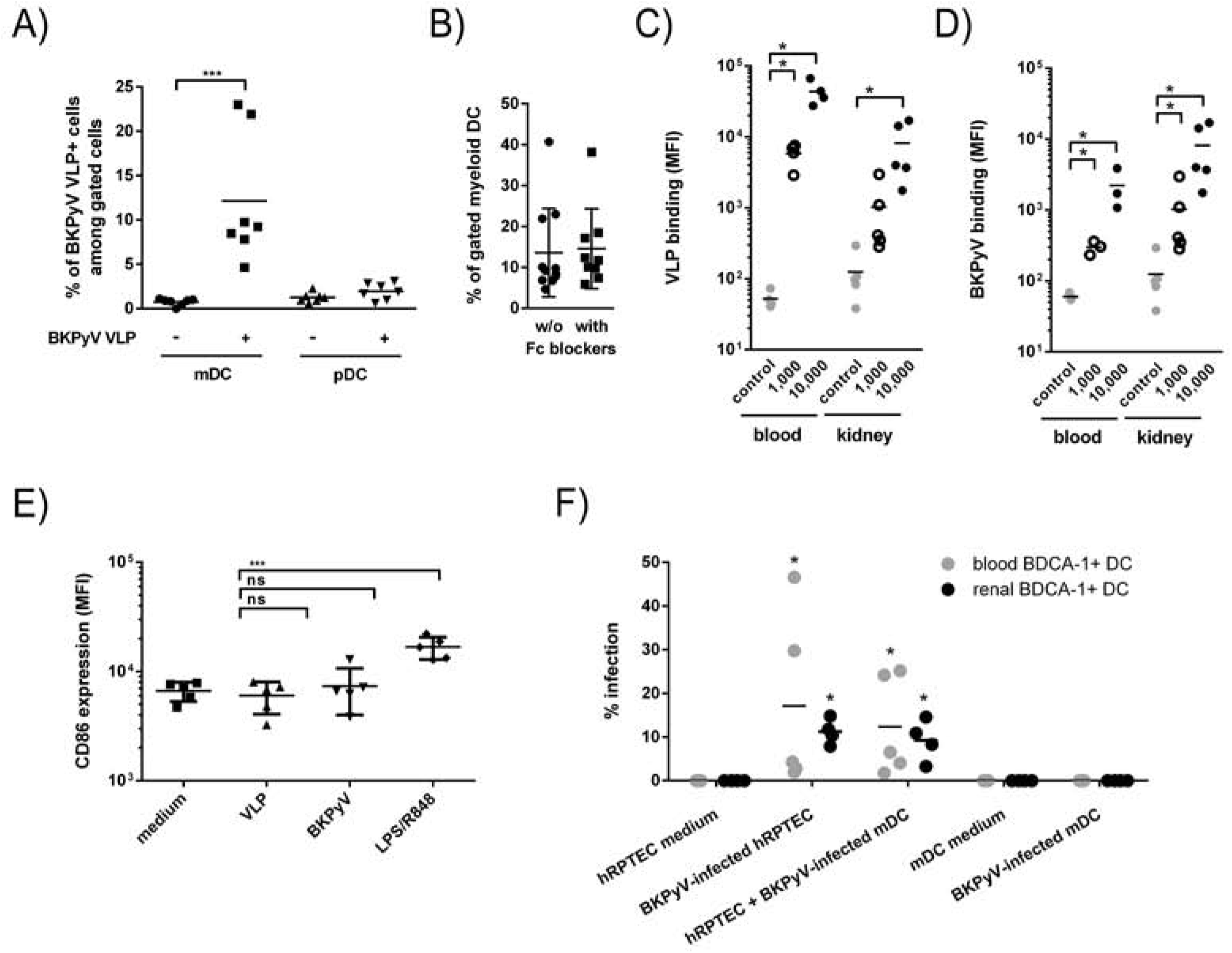
Blood and kidney CD1c^+^ myeloid DC bind and transmit BKPyV to primary hRPTEC without getting infected. (A) Quantitative measurement of VLP positive cDC and plasmacytoid (pDC) DC in whole blood of healthy volunteers according to the gating strategy shown in Supplemental Figure 1. (B) Quantitative analysis of VLP binding to cDC with (closed squares) or w/o (closed circles) Fc receptor blockade. Dose-dependent BKPyV VLP (C) or infectious particles (D) binding to purified CD1c^+^ cDC from blood or kidney of healthy individuals; the “control” condition means no VLP (grey circles). Black closed and empty circles represent 10^3^ and 10^4^ particles/cell respectively. The immunomagnetic cell sorting strategy is shown in Supplemental Figure 1. (E) CD86 cell surface expression assessed by flow cytometry on freshly isolated blood CD1c^+^ DC cultured for 24 hours in medium alone (squares) or with VLP (triangles; 103 particles per cells), BKPyV particles (inverted triangles; ibid) or with LPS/R848 (diamonds; 100ng/mL LPS and 1μg/mL R848); n=4 distinct blood donors. (F) Quantitative assessment of the ability of CD1c^+^ mDC from blood (grey dots) or kidney (black dots), ie from healthy blood donors and macroscopically healthy parts of resected human tumor-bearing kidneys, to capture and transfer BKPyV to hRPTEC as shown in Figure 6A. The percentage of infected hRPTEC corresponds to the percentage of LTAg+ hRPTEC within total cells, ie DAPI counterstained nuclei (42), based on an automated counting on a HCS device. Data are represented as percentage of infection. Statistical analyses have been applied to comparisons between % of infected hRPTEC in coculture with cDC with or w/o cDC preincubation with the Dunlop strain. Statistically significant results are marked by an asterisk; *=p<0.05, **=p<0.01, ***=p<0.001; one-way ANOVA with Tukey’s multiple comparison tests.

## Discussion

In this study, cDC, either generated *in vitro* or isolated from human blood and kidney, were shown to support BKPyV attachment in a sialic acid-dependent manner and subsequent clathrin-independent endocytosis through two distinct pathways, the first involving GRAF-1+ CLIC/GEEC and the second, minor pathway dependent on EEA1+ macropinocytic endocytosis. However, we did not provide evidence on potential spatio-temporal connections between both compartments. Upon contact, cDC were not activated by viral particles or BKPyV-infected cells. Moreover, we showed that MDDC and CD1c^+^ cDC were non-permissive to BKPyV infection. Internalized or membrane-bound BKPyV virions kept their ability to *trans*-infect permissive cells like hRPTEC and were also shown to be protected from neutralization by sera of BKPyV reactivating KTR.

We demonstrated that BKPyV interacts with human MDDC in a dose- and sialic acid-dependent manner suggesting these cells are equipped with BKPyV receptors, likely GD1b and GT1b(46). Upon attachment, BKPyV was shown to massively accumulate in pleiomorphous GRAF-1+ endocytic vesicles originating from the PM and partially overlapping with CTxB containing vesicles in MDDC. Although not proven here, it is tempting to speculate that a similar entry pathway for BKPyV occurs in CD1c-sorted cDC. This compartment was identified as CLIC/GEEC vesicles whose formation is clathrin-independent (see for review (72)). Interestingly, BKPyV was shown to infect hRPTEC in a clathrin- and caveolin-independent manner indicating that some BKPyV entry steps might be common between those cells and cDC(73, 74). Ewers and colleagues demonstrated that SV40 triggers the formation of PM invaginations related to CLIC/GEEC endocytosis after binding to GM1 in caveolin-1 deficient or energy-depleted cells(48). Multiple interactions between chemically defined GM1 PM clusters and SV40 capsomers were demonstrated to promote PM curvature and the formation of pleiomorphous tubules containing viral particles. CLIC/GEEC endocytosis has not been thoroughly documented for BKPyV before our results, even though Drachenberg et al observed BKPyV virions within tubular structures in hRPTEC from PVAN biopsies(30). Whether these tubular vesicles result from early endocytosis or from viral progeny remains to be clarified. Although the microtubule-associated motor protein, dynein 1, was shown not to play a role in BKPyV(75), JCPyV and SV40 infection(76), here we demonstrated that a specific chemical inhibitor of the dynein protein family caused a measurable reduction of the *trans*-infection process by MDDC indicating distinct cell-specific requirements for BKPyV entry. After internalization, BKPyV(77), JCPyV(77, 78), SV40(77, 79) and MuPyV(80) were shown to reach the ER within the first ten hours after cell attachment to hRPTEC(46, 75). This step is crucial for the infection(46, 75, 81, 82) since it is followed by the release of partially uncoated virions in the cytosol and import of viral genomes to the nucleus to initiate replication. This question was not directly addressed in the present study but the absence of LTAg in MDDC after several days pi strongly suggests either BKPyV does not undergo uncoating or that the CLIC/GEEC endocytosis does not lead to productive infection. In non-immune cells, CLIC/GEEC was shown as the main productive AAV2 infection route, invalidating the second possibility(83). These discrepancies between our results and former studies might reflect multiple common as well as distinct BKPyV entry steps according to the cell type studied. Further work is needed to establish the molecular determinants of such differences between non-immune and immune cells like cDC. A recent review pointed out the link between the CLIC/GEEC endocytosis and glycosphingolipids (GSL) which encompass gangliosides in the establishment of cell polarity(72). DC polarization leading to the formation of a synapse is an important event in T cell priming (see for review(84)) but might also be crucial in the BKPyV *trans*-infection process we described here. GSL are known to form lipid rafts on the PM (see for review(85)). Such micro domains function as a platform to segregate a wide range of effector molecules including GPI-anchored cargos(86). Wang and colleagues demonstrated that GPI-anchored molecules, which utilize the CLIC/GEEC endocytic pathway upon ligand binding, share biosynthetic pathways and common cellular locations with GSL(87). Such findings could link the ganglioside-mediated BKPyV attachment to viral endocytosis even in immune cells.

An important DC function is the ability to sense microbes through the recognition of conserved pathogen-associated molecular patterns (PAMP) by PM-bound (toll-like and C-type lectin receptors, TLR and CLR respectively), endosomal (TLR) or modified DNA/RNA cytoplasmic receptors altogether termed PRR(88). Viral particles as supramolecular arrangements of proteins and nucleic acids can be considered as PAMPs. Zepeda-Cervantes et al (Frontiers Immunol, 2020) have recently discussed numerous examples of VLP sensing leading to activation of human DC in a review(89). In contrast, both *in vitro*-generated murine and human DC were shown to remain unresponsive to BK- and JCPyV VLP(23, 26). Our results with human MDDC as well as freshly isolated blood and renal CD1c^+^ cDC confirm these observations and extend them to *bona fide* DC subsets supporting that such an immune ignorance towards BKPyV could exist *in vivo*. Two recent comprehensive studies demonstrated that hRPTEC fail to sense BKPyV(90, 91). This was shown to be partly dependent on the expression of the agnoprotein, a viral factor whose function has remained unclear so far(91). While BKPyV escape mechanisms seem to depend on viral gene expression in hRPTEC, we consider that different escape mechanisms are at work in BKPyV refractory cDC. The observed accumulation of BKPyV into CLIC/GEEC vesicles in MDDC after two hours might lead to their segregation in a PRR-free compartment. Unfortunately, whether PRR are present in the CLIC/GEEC compartment is unknown.

We demonstrated that although renal CD1c^+^ DC are refractory to BKPyV infection they remain able to capture virions and *trans*-infect hRPTEC *in vitro*. CD1c^+^ DC are normally present in the human renal interstitium surrounding the proximal tubules and glomeruli(29, 34, 71) but in PVAN lesions, a significant increase in infiltrating CD1c^+^ DC is documented(34). Whether the CD1c^+^ DC infiltrate has a key role in viral spreading *in vivo* deserves to be investigated through combined multidimensional imaging techniques and spatial RNA/DNA sequencing. Upon inflammation, monocytes are recruited in tissues where they differentiate in inflammatory DC with transcriptomic profiles closely related to those observed in MDDC(68, 69). PVAN develops in an inflammatory context. Therefore, it is tempting to speculate that along with resident CD1c^+^ DC, inflammatory DC could participate in the potentiation of BKPyV infection.

In this study, we demonstrated that cDC, namely MDDC and blood or kidney CD1c^+^ resident DC can capture infectious BKPyV through an unprecedented endocytic pathway in cDC and for BKPyV, and transfer the virus to permissive cells like hRPTEC without DC activation or infection, suggesting a role for cDC in BKPyV spreading. Moreover, we showed that internalized virions were protected from neutralization by serum from KTR. Taken together our results support the idea that cDC could facilitate BKPyV infection by favoring its spreading and limiting specific T lymphocyte activation due to the cDC ignorance towards BKPyV antigens and the circumvention of neutralization by specific antibodies. Hence, this work could help to understand how cDC could aggravate BKPyV infection in KTR.

## Supporting information

Supplemental Figues 1 and 2

## Author contributions

M.S. and F.H. designed, performed, analyzed and interpreted all experiments exept RNAseq data and wrote the paper. F.C., C.P., C.B., A.G., S.N., P.H., J.V., N.M., J.D., P.G., J.B.-G. performed and analyzed experiments. K.R., C.K.-A. and J.B. collected and characterized all human samples used in this study. A.T., R.J., D.McI. and C.B.-B. participated in experimental design and data discussion and actively contributed to the manuscript preparation. M.G. analyzed RNAseq data and deposited datasets on appropriate databases. F.H. supervised this study. All authors reviewed and approved the manuscript.

## Acknowledgments and financial disclosures

The authors would like to thank Camille Roesch (Izon Sciences Europe Ltd.) for her help in the VLP and BKPyV particle titrations based on the TRPS technique and Hélène Roux and Felix Letertre for their technical assistance. This work was supported by the Fondation d’Entreprises Progreffe (Mathieu Sikorski’s salary). This work was carried out in the context of the IHU-Cesti project, which received French government financial support managed by the Agence Nationale de la Recherche via the ‘‘Investment into the Future’’ program ANR-10-IBHU-005. The IHU-Cesti project is also supported by Nantes Metropole and the Pays de la Loire Region. The authors declare no conflict of interest. The funders had no role in study design, data collection and analysis, decision to publish, or preparation of the manuscript.

## Supplemental material

**Supplemental Figure 1:** Gating strategy of myeloid (CD11c+ in CD45+, HLA-DR+, Lin-cells) and plasmacytoid DC (CD123+ in CD45+, HLA-DR+, Lin-cells) in whole blood of healthy volunteers. Cells were incubated with 2.5μg/mL of Alexa Fluor®647 coupled-VLP or the same volume of PBS (no VLP; 45 minutes at 4°C) and representative dot plots showing the percentage of VLP+ cells.

**Supplemental Figure 2:** Assessment of purity of freshly isolated CD1c^+^ DC from blood and kidney. (A) Gating and enrichment evaluation before and after immunomagnetic cell sorting of CD19-CD1c^+^ myeloid blood DC. (B) Similar dot plots showing the purity of CD1c^+^ myeloid DC from resected human kidneys before and after the FACS-assisted sorting. These results are representative of five different cell isolations from both blood and kidney compartments.

