## Supplemental Figues 1 and 2 for "Non-permissive human conventional CD1c^+^ dendritic cells enable *trans*-infection of human primary renal tubular epithelial cells and protect BK polyomavirus from neutralization"

**Sikorski et al.**

### Supplemental material

**Supplemental Figure 1:** Gating strategy of myeloid (CD11c+ in CD45+, HLA-DR+, Lin- cells) and plasmacytoid DC (CD123+ in CD45+, HLA-DR+, Lin- cells) in whole blood of healthy volunteers. Cells were incubated with 2.5µg/mL of Alexa Fluor®647 coupled-VLP or the same volume of PBS (no VLP; 45 minutes at 4°C) and representative dot plots showing the percentage of VLP+ cells.

**Supplemental Figure 2:** Assessment of purity of freshly isolated CD1c+ DC from blood and kidney. (A) Gating and enrichment evaluation before and after immunomagnetic cell sorting of CD19- CD1c+ myeloid blood DC. (B) Similar dot plots showing the purity of CD1c+ myeloid DC from resected human kidneys before and after the FACS-assisted sorting. These results are representative of five different cell isolations from both blood and kidney compartments.

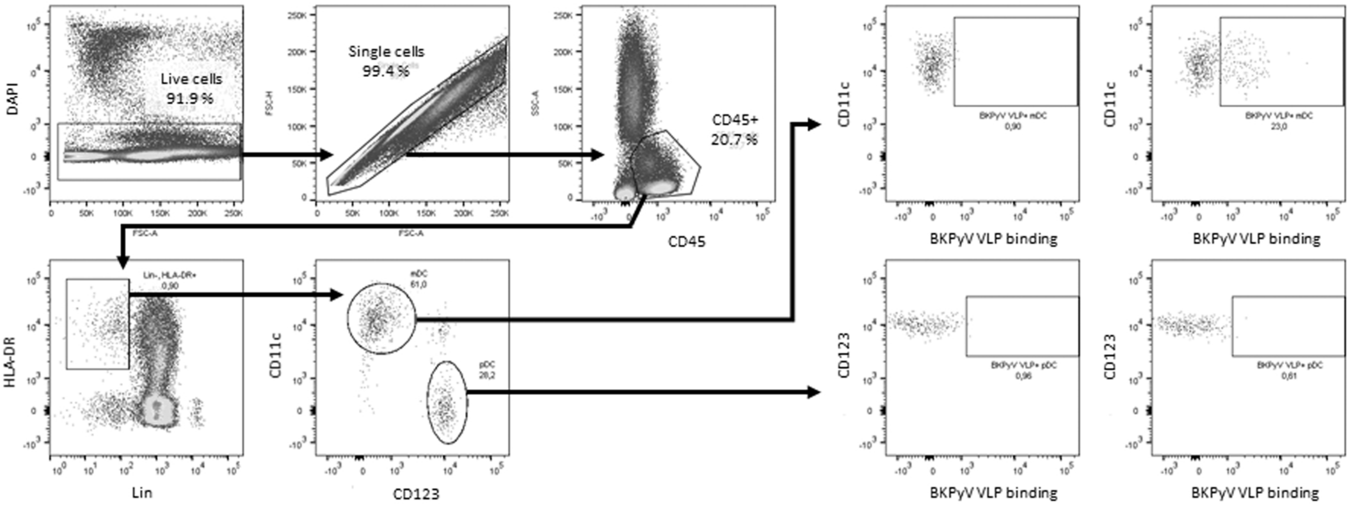


**Supplemental Figure 1: Gating strategy of myeloid (CD11c+ in CD45+, HLA-DR+, Lin- cells) and plasmacytoid DC (CD123+ in CD45+, HLA-DR+, Lin- cells)** in whole blood of healthy volunteers. Cells were incubated with 2.5µg/mL of Alexa Fluor®647 coupled-VLP or the same volume of PBS (no VLP; 45 minutes at 4°C) and representative dot plots showing the percentage of VLP+ cells.

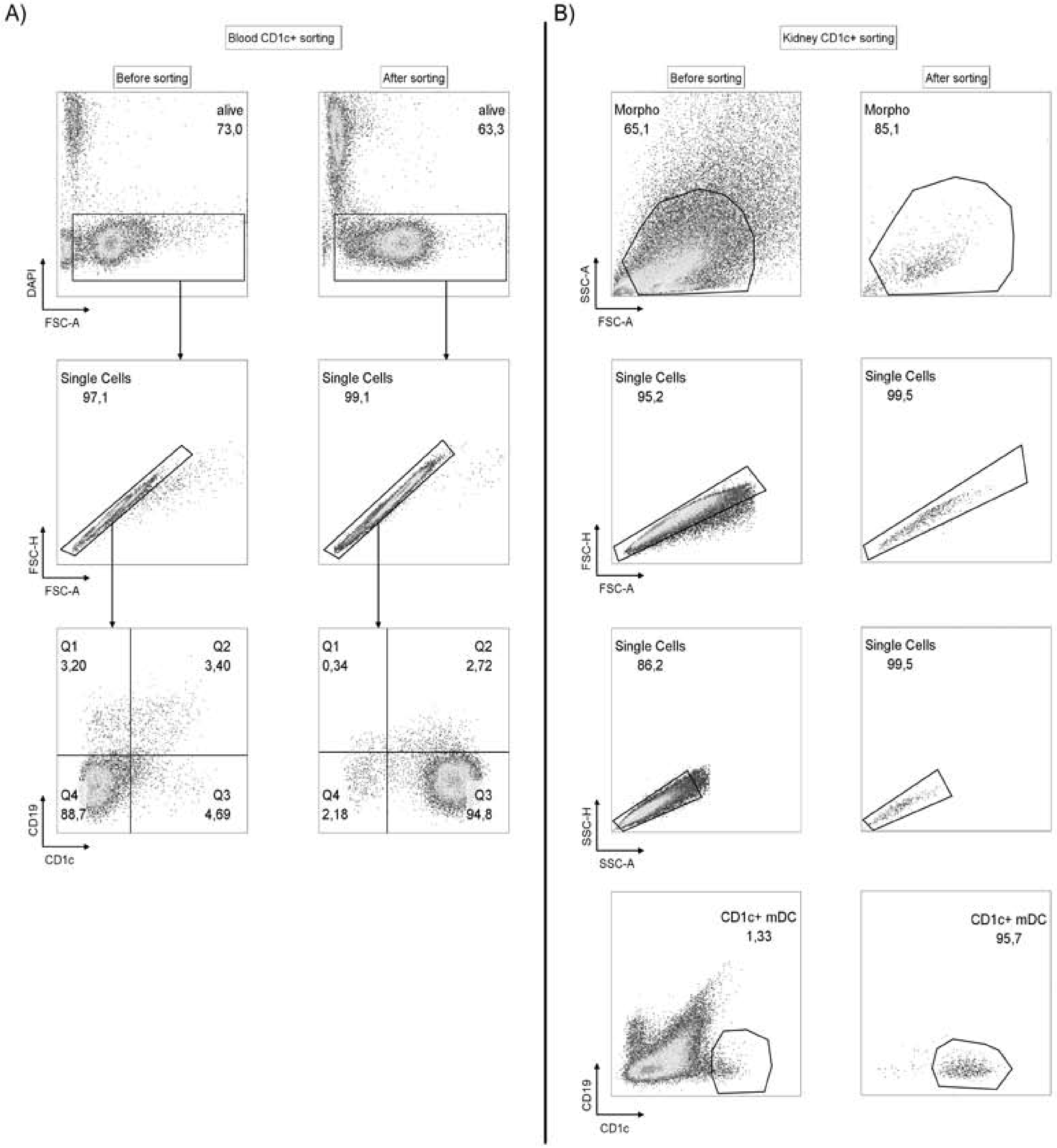


**Supplemental Figure 2: Purity of freshly isolated CD1c+ DC from blood and kidney.** (A) Gating and enrichment evaluation before and after immunomagnetic cell sorting of CD19- CD1c+ myeloid blood DC. (B) Similar dot plots showing the purity of CD1c+ myeloid DC from resected human kidneys before and after the FACS-assisted sorting. These results are representative of five different cell isolation from both blood and kidney compartments.
